# Data-Driven Strain Design Using Aggregated Adaptive Laboratory Evolution Mutational Data

**DOI:** 10.1101/2021.07.19.452699

**Authors:** Patrick V. Phaneuf, Daniel C. Zielinski, James T. Yurkovich, Josefin Johnsen, Richard Szubin, Lei Yang, Se Hyeuk Kim, Sebastian Schulz, Muyao Wu, Christopher Dalldorf, Emre Ozdemir, Bernhard O. Palsson, Adam M. Feist

**Author notes:** Corresponding Author: Adam M. Feist, University of California San Diego, 417 Powell-Focht Bioengineering Hall, 9500 Gilman Drive La Jolla, CA 92093-0412.

## Abstract

Microbes are being engineered for an increasingly large and diverse set of applications. However, the designing of microbial genomes remains challenging due to the general complexity of biological system. Adaptive Laboratory Evolution (ALE) leverages nature’s problem-solving processes to generate optimized genotypes currently inaccessible to rational methods. The large amount of public ALE data now represents a new opportunity for data-driven strain design. This study presents a novel and first of its kind meta-analysis workflow to derive data-driven strain designs from aggregate ALE mutational data using rich mutation annotations, statistical and structural biology methods. The mutational dataset consolidated and utilized in this study contained 63 *Escherichia coli* K-12 MG1655 based ALE experiments, described by 93 unique environmental conditions, 357 independent evolutions, and 13,957 observed mutations. High-level trends across the entire dataset were established and revealed that ALE-derived strain designs will largely be gene-centric, as opposed to non-coding, and a relatively small number of variants (approx. 4) can significantly alter cellular states and provide benefits which range from an increase in fitness to a complete necessity for survival. Three novel experimentally validated designs relevant to metabolic engineering applications are presented as use cases for the workflow. Specifically, these designs increased growth rates with glycerol as a carbon source through a point mutation to *glpK* and a truncation to *cyaA* or increased tolerance to toxic levels of isobutyric acid through a *pykF* truncation. These results demonstrate how strain designs can be extracted from aggregated ALE data to enhance strain design efforts.

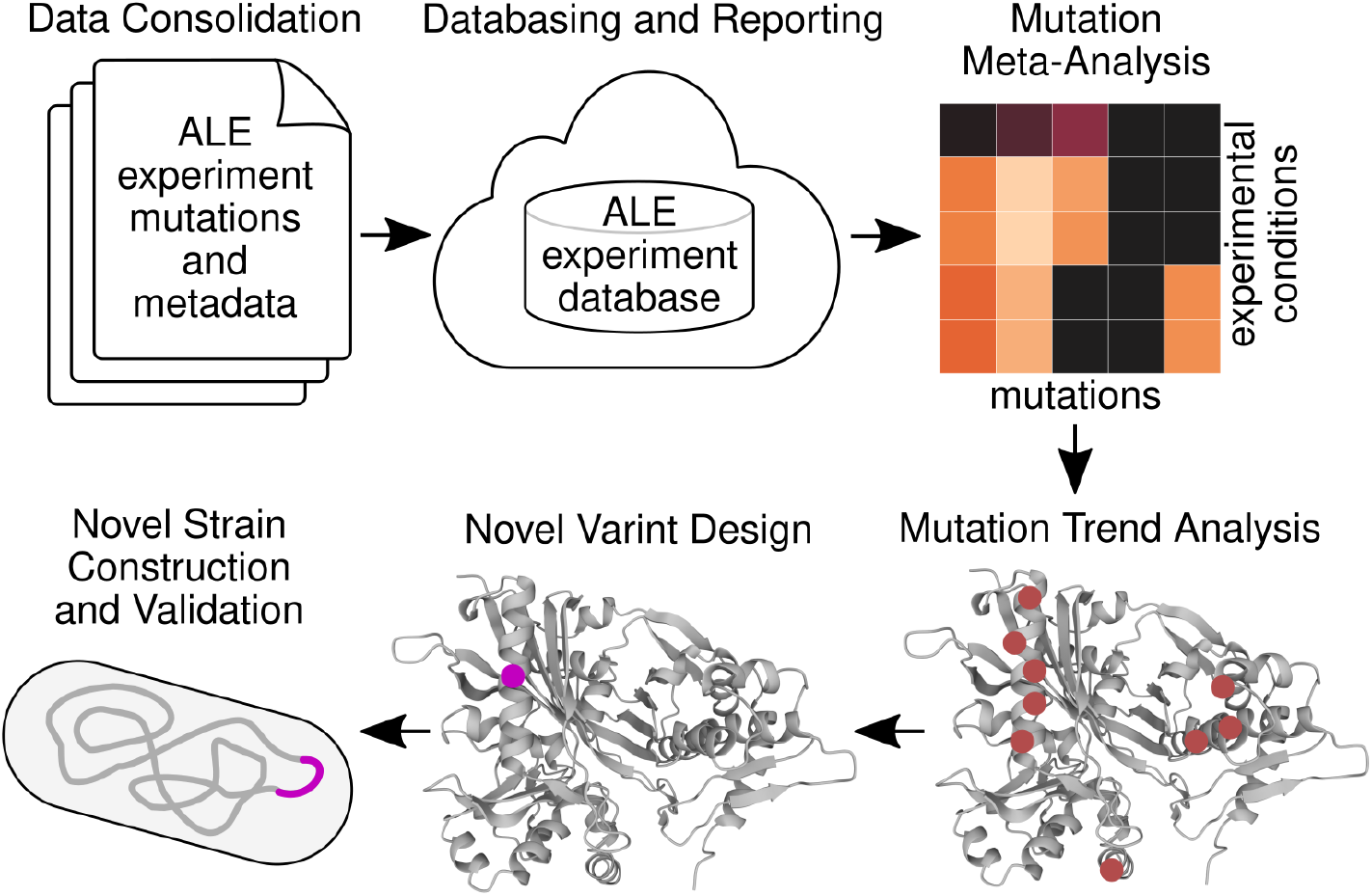

## 1 Introduction

The ability to precisely engineer microbial strains continues to improve. Multiple fields have emerged to take advantage of recent advances in biomolecular methods and technologies^1^. However, an incomplete understanding of biology renders the rational design of microbial strains challenging^2^. Introducing rationally designed changes into the DNA of a cellular chassis often leads to a perturbed suboptimal metabolic or regulatory state, resulting in the underachievement of goals^3^ and the need for a prolonged engineering effort that can require up to 6–8 years and over $50 million in costs^2^.

The opportunity exists to leverage the built-in problem-solving processes of adaptive evolution to discover and elucidate biological functions and generate solutions for applications. Adaptive Laboratory Evolution (ALE) is the formalization of a controlled evolutionary process that can successfully be applied to understand and engineer bacterial strains. ALE can provide adaptive mutations to optimize a strain’s growth rate or related fitness properties useful for both microbial engineering research and applications^4^. When a production pathway (e.g., metabolic) can be coupled with growth, ALE can provide adaptive mutations that will increase the throughput of this pathway^5,6^. Furthermore, if a strain’s fitness has been severely disrupted by a designed change, ALE can identify mutations that rebalance the cell’s homeostasis^4,7–11^. ALE can additionally serve to harden strains against industrial conditions^3,4,12,13^ and improve their utilization of secondary or non-native substrates^4,14–19^.

Due to ALE’s potential for discovery and application^4^, almost 700 manuscripts and over 18,000 experimental evolutions have been published^20^. The growth in ALE data has inspired multiple efforts towards its consolidation and analysis^20–23^. Aggregating public data is expected to enable new discoveries not evident through single experiments^24^. Critical biomedical efforts, such as cancer and antimicrobial resistance research, have benefited from the meta-analysis of big data sets describing their respective fields^25–27^. Previous work has datamined public ALE data, though their results were limited to genetic loci and therefore didn’t explore the specific mutational sequence changes^21^. Nucleotide-level resolution is ideal for strain design and the aggregation of ALE mutation sequence changes could reveal the specific types of changes adaptive evolution selects across defined genomic features and experimental conditions. These ALE-derived design principles could inform strain design efforts^28^. Thus, one could initiate a design based on an ALE mutation or evolved strain that best represents an apparent mutation trend from aggregated mutational data. One could further strive to understand ALE mutation trends well enough to propose novel sequence changes that would accomplish the same fitness benefit, demonstrating the possibility to extract strain design principles from ALE data.

This study seeks to design novel strain variants with potential for applications by using rich mutation annotations and structural biology methods on a consolidated ALE dataset of mutations and their experimental conditions. Aggregated ALE mutational and experimental conditions data was leveraged from ALEdb, a web-based platform reporting on experimental evolution mutations and their conditions^22^. The combination of rich annotations and aggregated ALE data enables mutated systems to be associated with experimental conditions to aid in deconvoluting mutation selection pressures^23^. Structural biology methods and functional annotations helped reveal the protein properties being targeted by ALE mutations. These analyses enable the interpretation of ALE mutation objectives for specific conditions of interest, which were leveraged to design and build novel variants of similar benefit to the ALE mutations. This work’s results demonstrate how to leverage existing aggregated ALE data and biological knowledge in microbial strain design and form a basis for similar applications in biomedical or basic discovery efforts.

## 2 Results

### 2.1 A workflow leveraging mutation trends in consolidated ALE data to design variants

A generalized workflow was developed to derive variant designs using aggregated ALE data (Figure 1). The following describes the workflow’s overall general steps and specific sections are referenced for each step to provide specific details:

**Figure 1:**
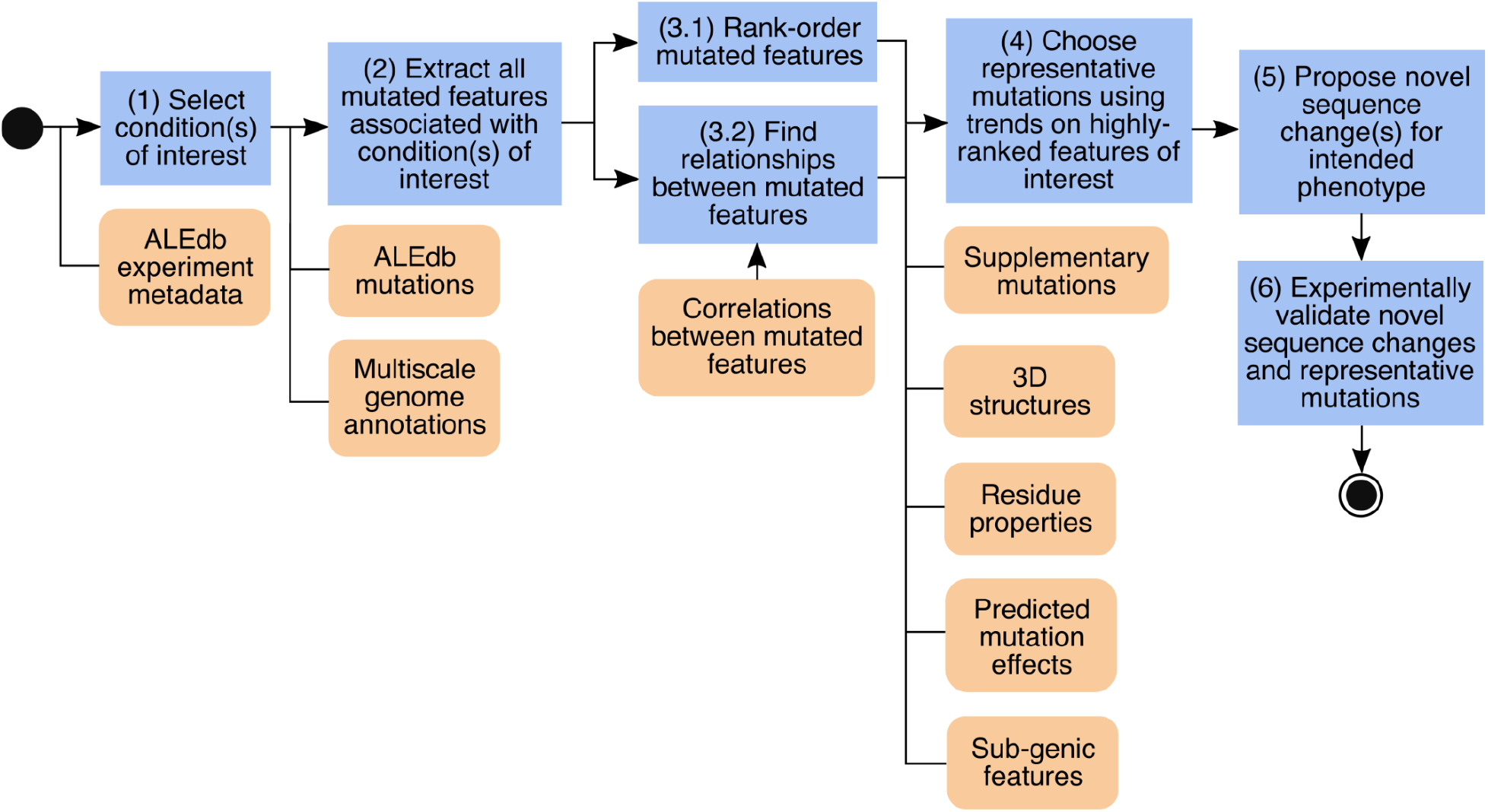
A general workflow to derive strain designs from aggregated ALE mutations and experimental conditions.

### 1. Select conditions of interest

This step is often dictated by the desired application and is informed by known biological mechanisms associated with the desired phenotype. The conditions available for investigation come from the metadata for ALE experiments which describe the experimental conditions (section 2.2). This study used ALEdb (7) as its source for ALE mutational data and experimental conditions. As an aggregated ALE dataset’s diversity increases, the queryable cases will also increase. In this work, growth on glycerol as a carbon source (section 2.3.1) and toxic concentrations of isobutyric acid (section 2.3.2) were targeted as relevant bioprocessing phenotypes^29–33^; there remains many other selection pressures in the dataset used.

### 2. Extract all mutated features associated with conditions of interest

Mutated genomic features and their experimental conditions are extracted from an aggregated and curated dataset to seed the analysis. Statistically significant associations are established between mutated features and conditions to aid in deconvoluting the selection pressures for mutated features. It is important to note that the scale of association analyses used to link mutated features to conditions depends on the amount and variety of annotated genomic features in a data set. Extending mutation annotations beyond the standard types included in genome references (genes and intergenic regions) improves the variety of mutated systems that may be associated with conditions of interest (section 2.2)^23^.

### 3. Rank-order the features of interest and identify relationships between them

Rank-order mutated features of interest by mutation frequency and strength of association to condition of interest (sections 2.3.1.1 and 2.3.2.1). Relationships between mutated features should also be considered (e.g., in the case of synergistic or antagonistic epistasis) and can help avoid potential incompatible sequence changes and provide insights into the systemic changes that result from multiple mutations (sections 2.3.1.1 and 2.3.2.1).

### 4. Choose representative mutations using trends on highly-ranked features of interest

Identify mutation trends on highly-ranked features of interest (sections 2.3.1.2, 2.3.1.5, and 2.3.2.2) and select representative mutations to establish initial ALE variant designs (sections 2.3.1.3, 2.3.1.7, and 2.3.2.3). Trends can be identified through mutation clustering on targets and can be informed using structural biology and mutation effect prediction methods.

### 5. Propose novel sequence changes

Interpret the ALE mutation trend’s objectives and propose a novel sequence change that would potentially accomplish the same fitness benefit as the representative ALE mutation. Interpretations can be informed using structural biology and mutation effect prediction methods, among others (sections 2.3.1.3, 2.3.1.8, and 2.3.2.3).

### 6. Experimentally validate novel sequence changes

Perform an assay to experimentally validate that the novel sequence changes are beneficial relative to wild-type and have similar fitness to ALE mutations (sections 2.3.1.4, 2.3.1.8, 2.3.2.4).

### 2.2 Meta-analysis trends suggest types of designs to expect from ALE data

This study specifically used ALE mutations and metadata from ALEdb, a database for experimental evolution mutation data^22^, and an enriched set of mutation annotations generated by a multiscale annotation method^23^. The dataset contained 63 *Escherichia coli* K-12 MG1655 based ALE experiments from ALEdb, totaling 357 independent evolutions, 13,957 observed mutations (Figure 2a). The observed mutations were filtered to exclude hypermutator strains, mutations with frequencies below 0.5 from population sequencing samples (24% of the total samples), and ALE-uniqueness, resulting in 3921 dominant ALE-unique mutations (Figure 2a). Mutations were annotated with 10 different genomic feature types (gene, intergenic, promoter, transcription factor binding site, ribosomal binding site, terminator, attenuator terminator, operon, pathway, regulon) to enable the identification of mutation convergence on a broad set of genomic features and biological functions^23^ (Figure 2a, Figure 3a). The dataset tracked 10 different condition types describing the strain and environment of ALE experiments with a total of 93 unique conditions (Figure 2a). Meta-analysis of this consolidated dataset revealed trends that predict the general shape of the workflow’s results and will be described in the following sections.

**Figure 2:**
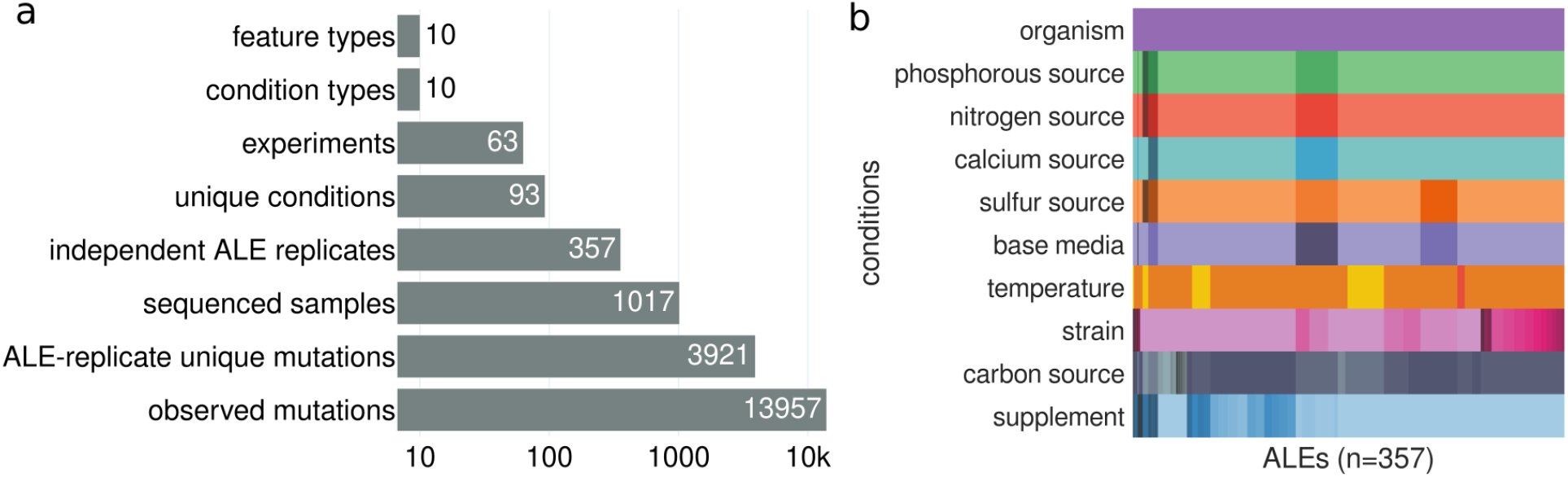
The dimensions and properties of the mutational data set used in this study. (a) A plot of the different dimensions of the ALE data used within this study as extracted from ALEdb. (b) A visual representation of the different condition types across ALEs in the targeted set. The mapping between individual colors and labels can be found in Supp. Fig 1-9.

**Figure 3:**
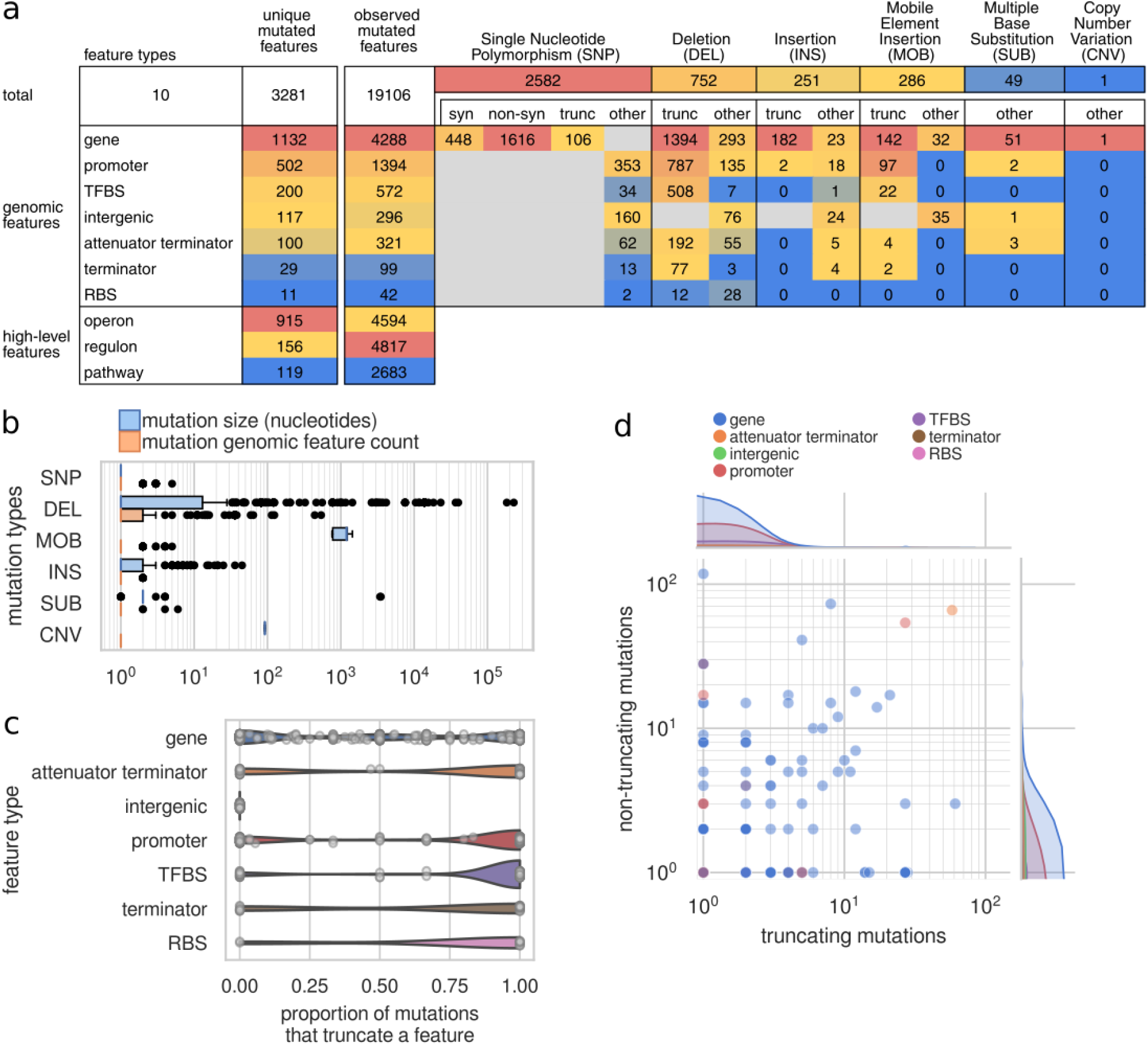
Mutation types and effects exhibit a bias towards specific genomic feature types. (a) Table of mutation type and mutated feature frequencies. Synonymous SNPs are abbreviated as “syn”, nonsynonymous SNPs are abbreviated as “non-syn”, and truncating mutations are abbreviated as “trunc”. (b) The distribution of mutation sizes and amount of genomic features affected according to mutation types. Abbreviations: SNP, single nucleotide polymorphism; DEL, deletion; MOB, mobile insertion elements; INS, insertion; SUB, substitution; CNV, copy number variant. (c) The proportion of mutations to individual features across feature types that are truncations. (d) The number of sequence truncating and non-truncating mutations for individual genomic features. Abbreviations: TFBS, transcription factor binding site; RBS, ribosomal binding site.

ALE mutations come in a variety of types and can affect a variety of features encoded on the genome. Six different types of mutations were found within the dataset (Figure 3a): single nucleotide polymorphisms (SNP), deletions (DEL), insertions (INS), mobile element insertions (MOB), multi-nucleotide substitutions (SUB), and copy number variations (CNV). Different mutation types manifest at different frequencies, resulting in different amounts for each mutation type within this dataset (Figure 3a). The mutated feature annotations used in this study range from small genomic features (e.g., terminators) to larger features (i.e., operons, regulons, and pathways) describing biological function (Figure 3a). There is a clear trend in the frequency of small mutated features: genes were most often mutated, with promoters hosting the most mutations for non-coding features. Multi-nucleotide mutations or overlapping features can result in more than one feature affected by a mutation, therefore more mutated features than mutations can occur within an ALE experiment^23^ (Figure 3a, Figure 3b).

#### 2.2.1 The majority of ALE mutations to noncoding regulatory features resulted in truncations

Amino acid substitutions and some of the disruptive effects mutations can have on genomic feature sequences can be confidently predicted. These predictions rely on mutation size (Figure 3b) or specific coding sequence changes. The effects of mutations on the translation of genes were the most straightforward to predict, primarily by considering the effects mutations have on the open reading frame. SNPs can result in synonymous or nonsynonymous codon changes, where a nonsynonymous SNP can result in an amino acid substitution or a truncation due to the introduction of a premature stop codon or the removal of a start codon. Truncations due to structural variants (SV) can also be predicted. Open reading frames were expected to be functionally truncated if the SV caused a frameshift. The effect of SV to non-coding regulatory features of the genome were more difficult to predict, though it is likely safe to assume that SVs of 10 nucleotides or more truncate both coding and non-coding features. SNPs were the most frequent mutation type and the largest contributor to mutated features (Figure 3a). SNPs were also most frequently found within genes and were most often predicted to result in a non-synonymous amino acid substitution, though the majority of these didn’t result in truncations (Figure 3a). Deletions were the most frequent SV and contribute a substantial amount of the truncations to features (Figure 3a).

Different feature types display different mutation type and effect trends. All non-coding feature types, besides intergenic regions, were more often targeted by truncations while genes were equally targeted by truncating and non-truncating mutations (Figure 3c). This suggested that non-coding features were more likely to be truncated than refined by ALE mutations. Individual features are often mutated with both truncating and non-truncating mutations, though there were features more often affected by one of the predicted mutation effects (Figure 3d). The features affected by only non-truncating mutations may be benefiting from gain-of-function mutations, which represent potential for variant designs involving something other than truncations.

#### 2.2.2 Unique mutated features are correlated with only a small number of other mutated features

Correlations between all mutated genomic features (gene, promoter, TFBS, intergenic, attenuator terminator, terminator, RBS) across the entire set of mutations can be leveraged to approximate coarse-grain relationships. Positively correlated genomic features should represent features that can be mutated together to optimize a strain for one or more conditions, which may constitute part of or the whole set of adaptive mutations from an ALE. Negative correlations should represent features that result in neutral or negative effects on the strain’s fitness when mutated together. The nature of selection for growth with ALE experiments results in more and stronger positive correlations than negative correlations between mutated features (Supplementary Figure 13, Figure 4a). Hierarchical clustering of correlated features results in a distribution of cluster sizes with a median of six features (Figure 4a, Figure 4b), potentially describing the general need for only six mutated features to achieve a system optimization through ALE.

**Figure 4.**
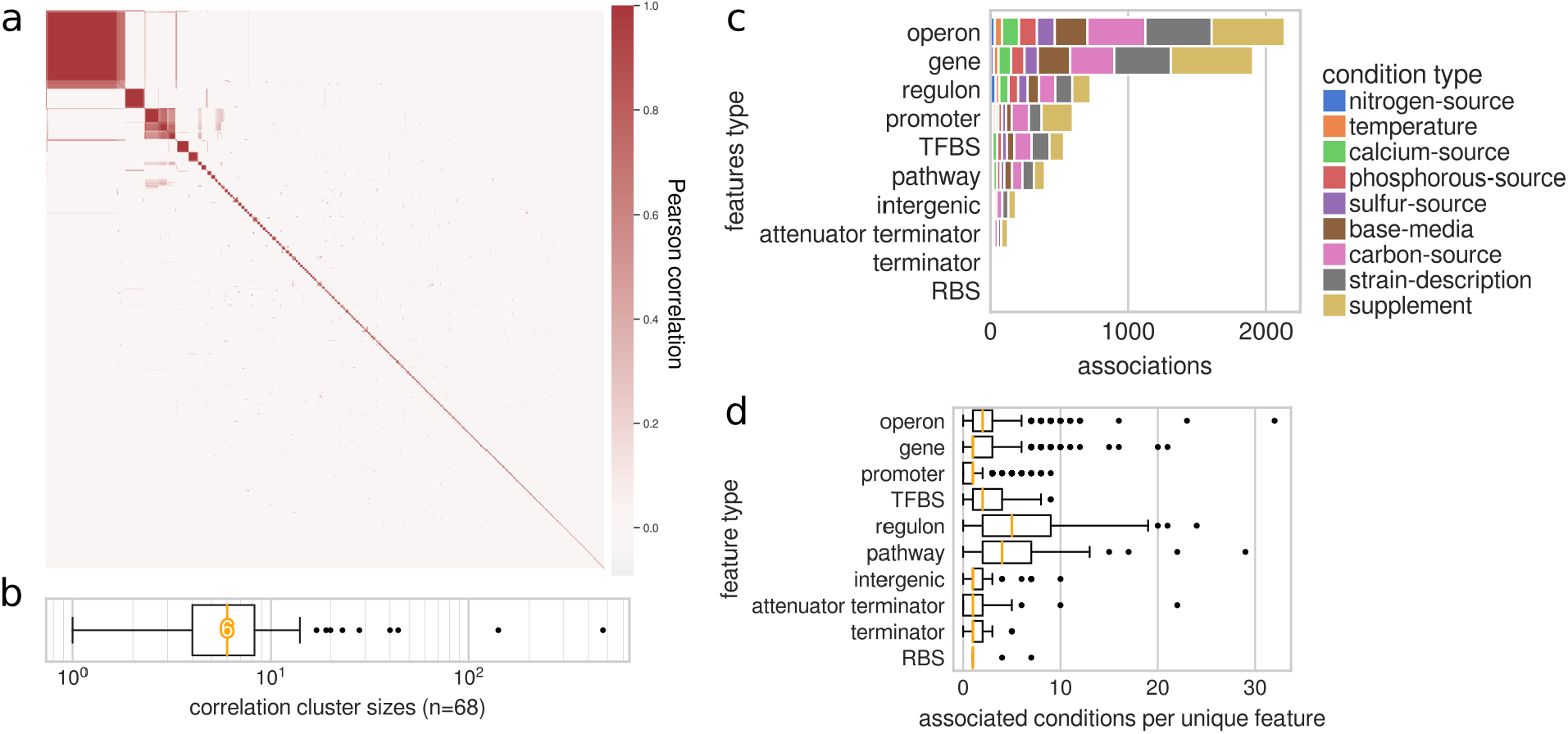
ALE adapted genotypes are gene-centric and involve few mutated features per condition. (a) A clustermap of the Pearson correlation coefficients for all genomic feature pairs (656,085). (b) The distribution of cluster sizes from the clustering of all genomic feature pairs according to their correlation. The median cluster size was 4, as highlighted. (c) The total amount of statistically significant associations between unique features and conditions according to feature types. (d) The amount of significantly associated conditions per unique genomic feature.

#### 2.2.3 Unique mutated features are associated with only a small number of conditions

The ALE conditions metadata enables efforts in linking mutated features to conditions, isolating subsets of mutations from across experiments that may be related to conditions of interest. There were 93 unique conditions across 10 different condition types (Figure 2b, Supplementary Figures 1 through 9). Due to the variety of experiments consolidated in the data of this study, some conditions will contain more potential for designs than others. The supplement, starting-strain, and carbon-source conditions have substantially more associated features than the other condition types (Supplementary Figure 11), likely reflecting the variety of conditions (Figure 2b, Supplementary Figures 1 through 9). Operons and genes had the largest amount of associated conditions (Figure 4c). This is likely due to operons being composed of genes, which have the largest variety of uniquely mutated features in this dataset (Figure 3a). These associations can also describe the specificity of the mutated features for the given conditions. Most features across all types were associated with only a small set of conditions, though there exist some outliers that were associated with a broad range of conditions and could therefore be applicable to a broad range of stresses (Figure 4d). According to these trends, it was expected that variants derived from these associations were going to be gene-centric and only involve a small number of coding and non-coding features.

#### 2.2.4 Aggregated ALE data reveals common low-frequency mutations targets across multiple experiments

Some mutated features are more often selected across independent ALE replicates of individual experiments and are described as having a measure of convergence^23^ (Figure 5, Supplementary Figure 10). The phenomenon of convergence is leveraged in evolution experiments to identify potentially beneficial mutations for given conditions. The degree of convergence that mutated features exhibit in their respective experiments provides a measure of selection strength for their mutation, suggesting an approximate degree of benefit for mutations. Mutated features generally have low convergence (Supplementary Figure 10), though there were some that demonstrate high convergence and therefore strong evidence of selection (Figure 5, Supplementary Figure 10). Those mutated features with high convergence are generally identified as hosting the most beneficial mutations within an ALE experiment. Some features mutated in many experiments were observed with low convergence and associated conditions (Figure 5), suggesting that these mutations were secondary optimizations beneficial to a broad set of conditions. These features may represent beneficial sequence changes that are only identifiable through the aggregation of ALE experiment mutations. For example, the *nagBAC-umpH* is mutated across 10 different ALE experiments in this dataset, though has an average convergence of 0.36 (Figure 5). *nagA* and *nagC*, the most frequently mutated genes of this operon, are involved in the recycling of cell wall peptidoglycan and may be introducing broadly applicable beneficial changes. Mutations to the cell envelope are often seen in ALE experiments and have been shown to be beneficial^34^.

**Figure 5.**
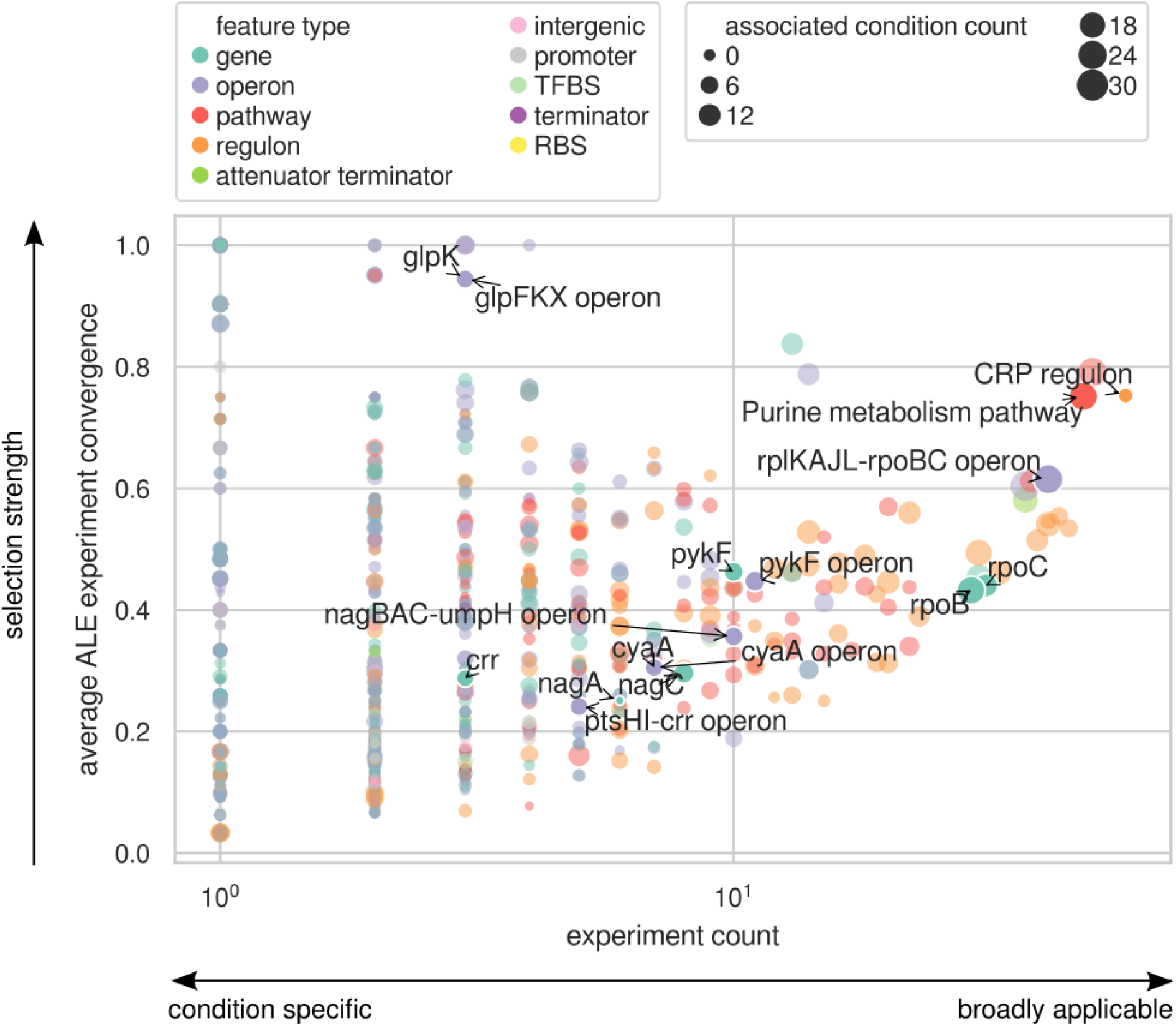
Aggregated ALE data reveals common low-frequency mutation targets with potential benefit to a broad set of conditions.

### 2.3 Case studies of application to strain design

The meta-analysis results reveal many opportunities for variant designs with the aggregated ALE dataset. The following sections describe case studies deriving three variant designs using ALE data and the presented workflow (Figure 1). The case studies demonstrate specific conditions that have a potential for application and have a large amount of samples or involve genes frequently mutated in multiple experiments. The case studies also demonstrate designs based on either truncating or non-truncating mutations to explore the potential for design of both mutation types.

#### 2.3.1 An *E. coli* K-12 MG1655 strain design for glycerol as a carbon source

The use of different carbon sources for bioproduction could prove to be an important strategy in maintaining feedstock flexibility. Glycerol has been shown to serve as feedstock for the production of valuable biochemicals, such as 1,3-PDO^29^, ethanol^30^, and limonene^31^. Glycerol-associated ALE mutations^23,35^ thus represent an opportunity for an ALE-derived strain design with possible valuable commercial applications and are targeted as a demonstrative case study (Figure 1 workflow step 1).

##### 2.3.1.1 Associations with conditions and analysis of multi-scale mutation annotations outlines systems important for adaptation to glycerol as a carbon source

Within the dataset, 149 mutations had at least one of their host features significantly associated with glycerol as a carbon source (Figure 1 workflow step 2, Supplementary Figure 14). The CRP regulon hosts the most mutated features for ALEs using glycerol as a carbon source (Supplementary Table 3), is associated with this selection pressure, and is one of the most frequently mutated regulons across this dataset’s ALE experiments (Figure 1 workflow step 2 and 3.1, Figure 5, Supplementary Figure 16). The CRP regulon describes many operons that encode for catabolic functions, including secondary carbon source metabolism. The CRP regulon is statistically associated with six different conditions, three of which were secondary carbon sources (Supplementary Figure 16). The three most frequently mutated operons associated with glycerol as a carbon source and linked to CRP were *glpFKX, cyaA*, and *ptsHI-crr* (Supplementary Table 2, Supplementary Figure 15, Supplementary Figure 16). The genes *glpK, cyaA*, and *crr* were the most frequently mutated features of their operons (Supplementary Figure 15, Figure 6a). Mutations to the CRP regulon, *glpFKX* operon, and *glpK* were strongly selected for (Figure 5) and *glpFKX* mutations were strongly associated with conditions involving glycerol and a temperature of 30 Celcius (Supplementary Table 1, Supplementary Figure 16). While the *cyaA* and *ptsHI-crr* operons and the *cyaA* and *crr* genes were less strongly selected for by their ALE experiments, they were still seen mutated in multiple ALE experiments (Figure 5) and were associated with a partially overlapping set of conditions (Supplementary Figure 16). Multiple strong associations to ALE conditions suggest the potential that mutations to these features could be beneficial for a broad set of stresses. Mutations to *glpK* and *cyaA* or *glpK* and *crr* were found in the same samples, though mutations to *crr* and *cyaA* were not (Figure 6a). This, along with correlations between the mutated genomic features associated with glycerol as a carbon source (Supplementary Figure 29), suggest that mutations to *crr* and *cyaA* have a negative epistatic relationship^23^ (Figure 1 workflow step 3.2).

**Figure 6.**
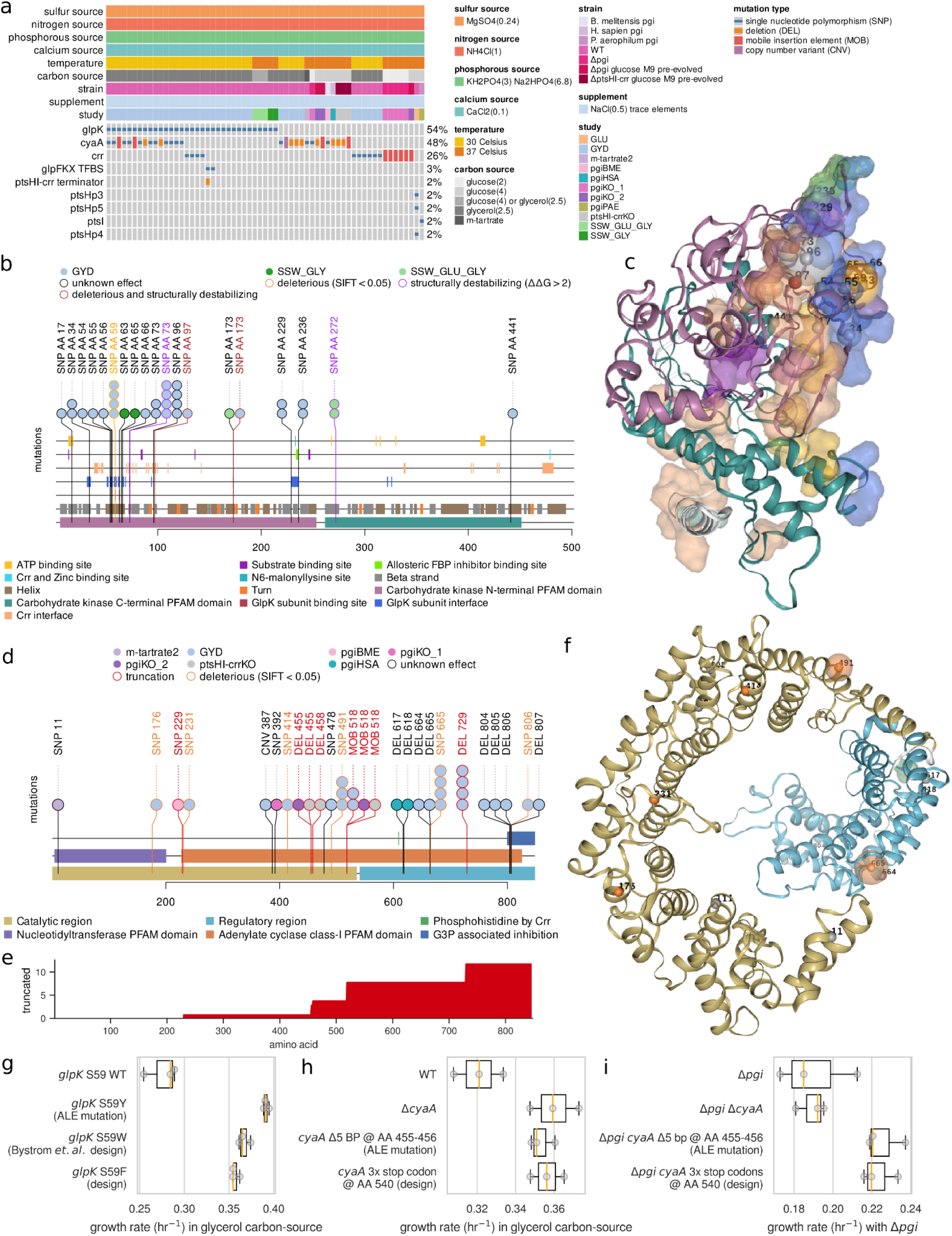
The clustering of truncating or non-truncating mutations reveal variant designs for glycerol as a carbon source. (a) An oncoplot demonstrating the types of mutations to genomic features on operons of interest (operons *cyaA, glpFKX*, and *ptsHI-crr*) and the conditions for the ALE samples hosting these mutations. (b) A mutation needle plot for mutated amino acids across GlpK’s amino acid chain. (c) GlpK’s 3D structure and mutated residues from mutations. The residue chain and transparent surfaces are colored according to the legend of the corresponding mutation needle plot. Mutations are represented by a small opaque sphere with a value representing their amino acid position on the corresponding mutation needle plot. The color of the mutation’s sphere corresponds to the mutation’s predicted effect as described by the legend on the corresponding mutation need plot. The transparent sphere centered on the mutations’ opaque sphere represents the number of mutations with a specific predicted effect on that position. The angle shown illustrates how all the GlpK-GlpK interface surfaces are oriented on the same side of the 3D structure along with the clustering of mutations on or near these surfaces. (d) A mutation needle plot for mutated amino acids across CyaA’s amino acid chain.(e) The accumulation of the truncated amino acids downstream of truncating mutation from the mutation needle plot. (f) CyaA’s protein structure and mutated residues from non-truncating mutations. (g) The growth rates of the mutants harboring ALE mutations and designed variants for GlpK in the selection pressure of glycerol as a carbon source. (h) The growth rates of the mutants harboring ALE mutations and designed variants for CyaA in the selection pressure of glycerol as a carbon source. (i) The growth rates of the mutants harboring ALE mutations and designed variants for CyaA in the selection pressure of Δ*pgi*.

##### 2.3.1.2 ALE mutation trends in *glpK, cyaA*, and *crr*

Among *glpK, cyaA*, and *crr, glpK* was the most frequently mutated (Figure 1 workflow step 3.1). Further, it was observed to mutate in ALE experiments that involve substrate switching between glucose and glycerol (Figure 6a); all ALE experiment mutations to *glpK* were investigated to understand if mutation types clustered within the gene according to conditions. All mutations to *glpK* were SNPs and frequently landed in codons for amino acids involved in GlpK’s subunit interface (Figure 6b, Figure 1 workflow step 4), where GlpK can form both a dimer and tetramer. The subunit interface amino acids are spread across GlpK’s sequence (Figure 6b), though in the 3D structure model their residue surfaces all group together to face the same direction (Figure 6c), revealing further clustering of mutations specific to 3D space. Finding the distances between mutated residues and GlpK features according to their 3D positions on GlpK’s structure provided for a potentially more accurate measure of nearness between mutations and features. GlpK subunit interfaces continue to be nearest to or directly host the most SNPs (Supplementary Figure 17), though the mutations to the GlpK subunit binding sites have the highest proportion of mutations with a predicted effect (Supplementary Figure 18). Four out of five of the SNPs to the GlpK subunit binding sites were accomplished through the same nucleotide substitution S59Y (TCC→ TAC) and was predicted deleterious via a SIFT score < 0.05.

*cyaA* was the second most frequently mutated gene of those associated with glycerol and its mutations were often found in samples with mutations to *glpK* (Figure 6a). cyaA was mutated in multiple experiments of different conditions (Figure 6a); all ALE experiment mutations to cyaA were investigated to understand if mutation types clustered within the gene according to conditions. Mutations from multiple experiments cluster near the middle of the amino acid sequence (Figure 6d, Figure 1 workflow step 4). Many of the mutations within this cluster cause a frameshift or truncation, thereby disrupting the downstream coding sequence. The accumulation of truncated amino acids demonstrated that the regulatory region was often targeted for disruption across varying ALE stressors, including that of glycerol as a carbon source (Figure 6e). Observing the mutated residues on CyaA’s structure from mutations other than truncations, there was no novel 3D clustering of mutated residues not already apparent with the linear analysis (Figure 6f). The clustering of mutations from multiple ALE experiments with various selection pressures suggests that a specific change to *cyaA* may have broad applicability.

*crr* was the third most frequently mutated gene of those associated with glycerol (Figure 6a) and its mutations were often found in samples with mutations to *glpK* and never found in samples with mutations to *cyaA* (Figure 6a, Figure 1 workflow step 3.2). The absence of mutated *cyaA* and *crr* genes in the same sample has been hypothesized to be due to their mutations having similar effects to the same system in the case of ALEs with a glycerol carbon source^23^. *crr* was mutated in multiple experiments of different conditions (Figure 6a); all ALE experiment mutations to *crr* were investigated to understand if mutation types clustered within the gene according to selection pressure. Most mutations to *crr*’s amino-acid sequence fall on or near interfaces (Figure 1 workflow step 4). These interfaces describe separate surfaces of Crr’s structure^36^ (Supplementary Figure 21). Finding the distances between mutated residues and Crr features according to their 3D positions on Crr’s structure^36^ provided for a potentially more accurate measure of nearness between mutations and features. Crr feature residues truncated by an upstream coding disruption were additionally included. The Crr-Crr interface hosts the most mutated residues (Supplementary Figure 22), though its mutation type differs from those mutated residues to all other features (Supplementary Figure 23). Mutations to the Crr-Crr interface were also specific to Δ*pgi* ALEs while mutations to all other features were specific to glycerol carbon source ALEs (Supplementary Figure 20). Mutations from glycerol carbon source ALEs additionally cluster near each other on or near the same surface of the 3D structure (Supplementary Figure 21). This surface hosts the GlpK, PtsG, PtsH, PtsI, and FrsA interfaces as well as binding and active sites. All but one mutation clustering near these multi-interface residues were predicted to be disruptive (Supplementary Figure 20).

##### 2.3.1.3 *glpK* and *cyaA* novel variant designs

The S59Y (TCC→ TAC) substitution was chosen as the representative ALE mutation for *glpK* due to its high frequency and specificity of effect on an active site (Figure 1 workflow step 4). Across all possible amino acid substitutions for *glpK* S59, the tyrosine substitution (Y) was very highly ranked according to SIFT scores, size, and flexibility difference relative to wild-type residues (Supplementary Figure 19). The residue substitution of tryptophan (W) scored higher or similar to the tyrosine ALE-derived substitution in the previously mentioned categories, as well as being categorized as having a bulky aromatic sidechain (Supplementary Figure 19b), thus making it a good candidate for a derived design. A substitution of tryptophan at this position has already been characterized to eliminate inhibition of GlpK’s catalytic activity^37^. Since the tryptophan substitution had already been characterized, another novel substitution was pursued for the purposes of this study. Phenylalanine (F) was the only other residue that is both characterized as being bulky and having an aromatic sidechain as well as scoring similarly to tyrosine and tryptophan substitutions; therefore, the final proposed design for GlpK was that of S59F (Figure 1 workflow step 5).

Mutations to *cyaA* have a more interpretable effect on the gene than those to Crr: the truncation of CyaA’s regulatory region. The 5 BP deletion starting within amino acid 455 in *cyaA* was chosen as the representative ALE mutation for those of CyaA and Crr (Figure 1 workflow step 4). This mutation was selected due to being mutated in two separate ALE replicates along with being an easily reintroducible mutation type. To truncate the regulatory region with more accuracy, CyaA’s variant design inserted 3 stop codons at amino acid 540, immediately upstream of the regulatory region (Figure 1 workflow step 5).

##### 2.3.1.4 Experimental validation of *glpK* and *cyaA* novel variant designs

To examine the fitness changes from *glpK* and *cyaA* mutants relative to wild-type with glycerol as a carbon source, growth screens were performed on reconstructed strains harboring the ALE mutations with the designed variants. The results show that the ALE-derived mutants and designs have similar growth rates along with higher growth rates than wild-type (Figure 6g, Figure 6h, Figure 1 workflow step 6).

To test the potential applicability of ALE-data-driven-designs with multiple different stresses, the *cyaA* mutants and design were additionally tested in the background of a Δ*pgi* strain. Some of the *cyaA* ALE mutations that clustered in the center of the gene were selected for by a Δ*pgi* ALE experiment (Figure 6d). The growth screen results demonstrated that the mutants and designs have similar growth rates along with higher growth rates than wild-type (Figure 6i). These results also show that a partial truncation to *cyaA* granted a higher fitness than a full truncation (Figure 6i, Figure 1 workflow step 6), which differs from the glycerol carbon-source stressor growth screen, where partial and full truncations had similar fitness.

#### 2.3.2 An *E. coli* K-12 MG1655 strain design for high concentrations of isobutyric acid

Tolerance is a key phenotype for microbial cell factories. Industrial requirements can have strains exposed to or produce toxic concentrations of substrates or products. Genotypes that provide any improved tolerance to detrimental conditions can be valuable in that their tolerance can translate to higher concentrations of product, especially with large scale operations^32^. isobutyric acid is a biochemical with a market size of 100,000 tons in 2011^33^ and can be produced with an engineered *E. coli* strain^32^. isobutyric acid associated ALE mutations^32^represent an opportunity for an ALE-derived strain design with possible applications (Figure 1 workflow step 1).

##### 2.3.2.1 Associations with conditions and analysis of multi-scale mutation annotations outlines systems important for adaptation to isobutyric acid tolerance

Within the dataset, 79 mutations had at least one of their host features significantly associated with the condition of toxic isobutyric acid concentrations (Supplementary Figure 24, Figure 1 workflow step 2). The purine metabolism pathway hosts the most mutations (Supplementary Table 6), with the majority of the mutations coming from the *pykF* and *rplKAJL-rpoBC* operons (Supplementary Figure 25, Supplementary Table 5, Figure 1 workflow step 3.1). Of these two, the *pykF* operon is strongly associated with toxic isobutyric acid concentrations (Supplementary Figure 28). The purine metabolic pathway, *rplKAJL-rpoBC* operon, *pykF* operon, and *pykF* were frequently mutated features across this dataset’s ALE experiments (Figure 5). The purine metabolic pathway and *rplKAJL-rpoBC* operon were strongly selected for in ALE experiments, while mutations to *pykF* and its operon were less so, on average (Figure 5). *pykF* and its operon were also more specific in their associations to conditions, where the *rplKAJL-rpoBC* operon and purine metabolic pathway mutation associations were more broad (Figure 5, Supplementary Figure 28). Correlations between frequently mutated genomic features demonstrate *pykF* and *rpoB* as positively correlated (Supplementary Table 4, Supplementary Figure 30). The positive correlation between *pykF* and *rpoB* along with their differing associations indicate that these compatible mutations were adapting for different selection pressures (Figure 1 workflow step 3.2).

##### 2.3.2.2 ALE mutation trends in *pykF*

The *pykF* operon is frequently mutated across this study’s many ALE experiments. These mutations were gathered and investigated to understand if mutations clustered according to ALE experiment conditions. Most of these mutations target *pykF’s* coding sequence, with some to the operon’s non-coding features (Figure 7a). A mutation to the *pykF* operon’s non-coding features seems to preclude a mutation to any of the others, suggesting that they have related effects on the host strain (Figure 7a). Mutations to *pykF* were spread across its sequence, with the majority contained in the second half of the sequence (Figure 7b). Due to the large variety of experiments mutating *pykF*, the broad distribution of mutations across *pykF*’s sequence, and the large amount and variety of structural feature annotations available for PykF, it is expected that the trends described by the aggregate of all ALE experiment mutations would be most revealing. Many of the mutations to *pykF* were predicted to truncate the coding sequence or disrupt the potential function and/or structural stability of a domain through a non-synonymous amino acid substitution. Considering the accumulation of truncated amino acids, the PykF subunit interfaces were the most frequently mutated features on PykF (Figure 7b, Figure 7d, Supplementary Figure 26). The truncations also seem to cluster on the second half of PykF’s sequence, resulting in the downstream coding disruptions primarily affecting the PykF tetramer subunit interfaces while avoiding the active sites in the first half of the sequence (Figure 7b, Figure 7d). Those mutations predicted not to cause truncations or frameshifts were found on PykF’s 3D structure nearest to binding sites within the cleft of PykF’s barrel domain (Figure 7c, Supplementary Figure 27). All mutations to PykF and those specifically from the toxic isobutyric acid ALEs follow the same trend across features on PykF’s structure (Supplementary Figure 26), providing evidence that mutations to *pykF* across different sets of conditions may accomplish similar outcomes. These trends suggest unique functional targets for the different mutation types, with truncations having the clearest outcome: a disruption to PykF’s ability to form a complex (Figure 1 workflow step 4).

**Figure 7.**
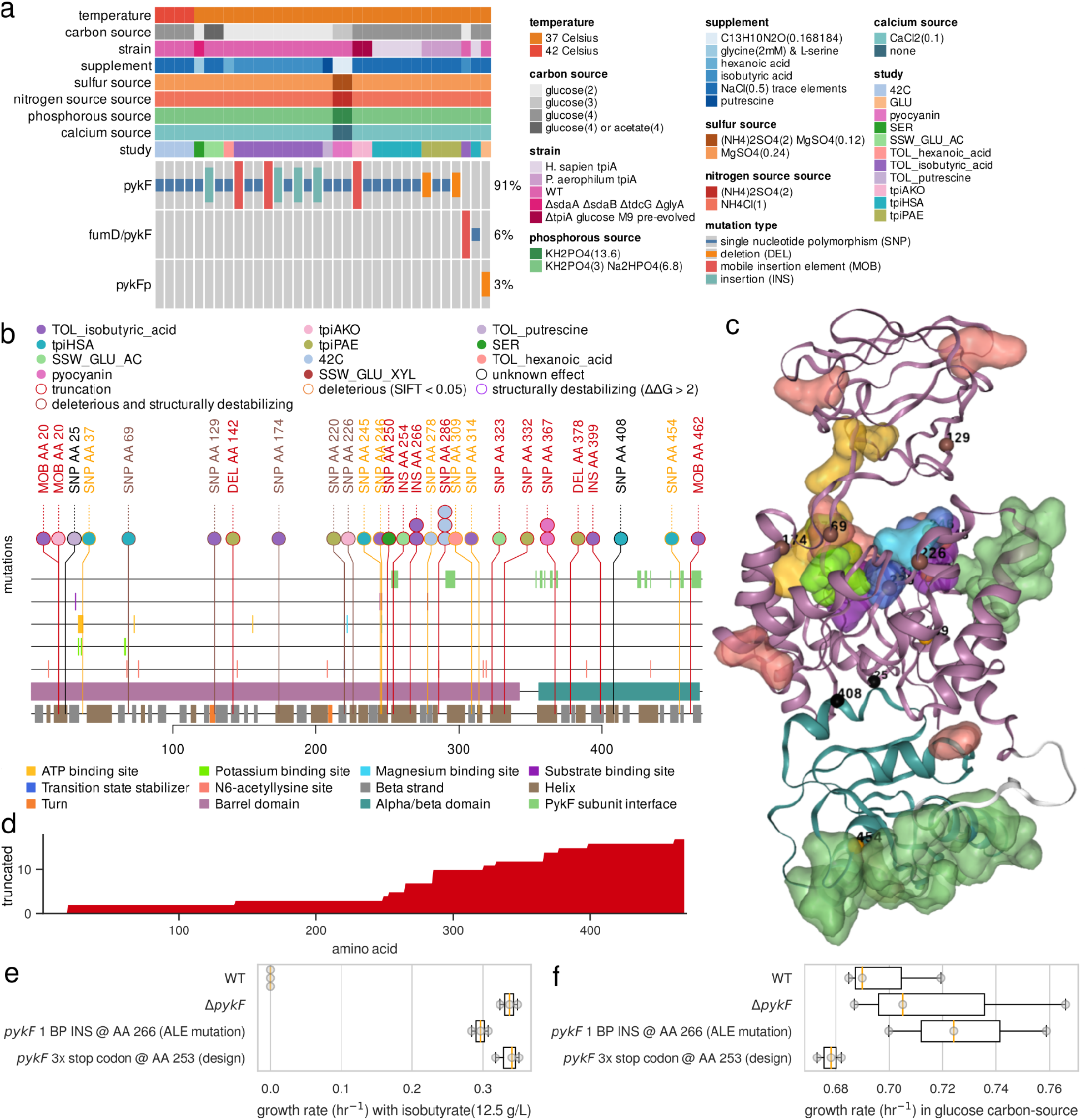
The clustering of truncating mutations reveals a variant design for toxic concentrations of isobutyric acid. (a) An oncoplot demonstrating mutations linked to the *pykF* operon across all ALE experiments of this study’s data. (b) A mutation needle plot for mutated amino acids across PykF’s amino acid chain. (c) PykF’s 3D structure and mutated residues. No truncating mutations are included. The residue chain and transparent surfaces are colored according to the legend of the corresponding mutation needle plot. Mutations are represented by a small opaque sphere with a value representing their amino acid position on the corresponding mutation needle plot. The color of the mutation’s sphere corresponds to the mutation’s predicted effect as described by the legend on the corresponding mutation need plot. The transparent sphere centered on the mutation’s opaque sphere represents the number of mutations with a specific predicted effect on that position. The angle shown illustrates how most of the mutations cluster in 3D space around the area which hosts most of the catalytic domains. (d) The accumulation of the truncated amino acids downstream of truncating mutations from mutation needle plot. (e) The growth rates of WT, a *ΔpykF* strain, the *pykF* ALE mutant, and the *pykF* designed variant with toxic concentration of isobutyric acid (12.5 g/L). A *ΔpykF* mutant was used to investigate for any difference between the strains that partially truncate *pykF* and its full truncation. (f) The growth rates of WT, a *ΔpykF* strain, the *pykF* ALE mutant, and the *pykF* designed variant with glucose as a carbon source. A *ΔpykF* mutant was used to investigate for any difference between the strains that partially truncate *pykF* and its full truncation.

##### 2.3.2.3 *pykF* novel variant design

A truncating insertion to amino acid 266 was chosen as the representative ALE mutation to *pykF* for the toxic isobutyric acid condition since it represents a clear trend in disrupting the PykF subunit interfaces and manifested in two independent ALE replicates with this selection pressure (Figure 1 workflow step 4). The interpretable trend of truncated amino acids accumulating across PykF subunit interfaces led to a variant design that truncated all of the PykF subunit interfaces: an insertion of 3 stop codons at amino acid 253, immediately upstream of all the PykF subunit interfaces (Figure 1 workflow step 5). These ALE mutations and the derived genome design were initially inspired by their strong selection with the isobutyric acid tolerance ALEs, though their benefit may instead be directly derived from growth on an abundance of glucose as a carbon source. All ALEs with mutations to *pykF* include abundant glucose as a carbon source (Figure 7a).

##### 2.3.2.4 Experimental validation of *pykF* novel variant design

Growth screens were performed on the reconstructed strains harboring the *pykF* ALE mutation along with the designed truncation to examine the differences in fitness between mutants and wild-type (Figure 1 workflow step 6). Growth screens for both toxic isobutyric acid levels and competitive glucose uptake were performed. The results show the mutant and designs have similar growth rates as well as higher growth rates than wild type with toxic isobutyric acid concentrations (Figure 7e) while not demonstrating obvious benefit in competitive glucose screens (Figure 7f).

## 3. Discussion

The phenomena of convergent mutations with experimental evolution has served well to reveal genotypic changes that likely result in a higher level of fitness for the host strain. Experimental evolution methods such as ALE leverage convergence to hypothesize which mutations bring about fitness benefits. ALE has been successfully applied in strain engineering, though the resulting mutation sets can be too small to deduce the functions in which the ALE mutations are targeting. This work demonstrates how aggregated public ALE mutation data, rich annotations, and structural biology methods can provide sufficient evidence to interpret ALE mutation objectives towards strain design.

Multiple methods were implemented to find ALE mutation trends at different levels of detail. These methods were organized into a workflow so that the sequential execution of steps leads to the design of sequence variants. Each step of the workflow was critical to deriving variant designs since individual steps worked to narrow down the ALE mutation targets and objectives at different levels of detail. Associations between mutated features and conditions extracted the mutated features potentially relevant for a condition of interest with strain design. Multi-scale annotations helped group mutated features that belong to the same system, clarifying whether a beneficial change can be accomplished through few or many mutated features. A meta-analysis of the new mutational dimensions resulting from these methods offers insights for variant designs derived from ALE mutational data. The majority of ALE mutations to non-coding regulatory features resulted in predicted truncations. Biological robustness is achieved by the regulation and maintenance of a variety of biological functions. Deregulation of biological functions to remove restrictions on flux or the deactivation of unnecessary functions may ultimately benefit a host’s performance in highly specific ALE environmental selection pressures. In essence, the optimization of an organism through ALE may result in a simplification of its systems towards maximizing the use of a subset of beneficial functions. Most ALE-data-derived genotypic solutions for a selection pressures may only require a small number of sequence changes and those sequence changes will most likely be in genes. This was shown by the small median cluster size of correlated mutant genomic features and the small median amount of conditions associated with mutated features. Genes were, by far, the most frequently mutated genomic feature. The bias towards genetic mutations may be due to many factors, one of which is that the genome of *E. coli* K-12 MG1655 is mostly composed of coding sequences (24). Another potential factor is that this study’s ALE selection pressures may only have required small alterations to functions encoded by genes. Finally, the meta-analysis and case-study results demonstrated that the same mutated gene or the exact same variant can provide fitness across a variety of stressors. This result emphasizes the value of aggregated ALE data in that it enables the identification of broadly applicable variants.

The case studies in this work demonstrate that enriched ALE data and analysis methods as organized in the strain design workflow provided enough evidence to deduce the sequence change properties and resulting molecular mechanisms converged upon by ALE mutations. From these results, sequence-based variants were derived and revealed insights into ALE-data-driven strain design. Two different variant design types were derived from two different mutation trends. The variant designs involving truncations were interpreted from the mutation trend of accumulating truncated amino acids on functional annotations across a gene’s amino acid sequence. The variant designs involving non-truncating mutations were interpreted from the clustering of mutated residues on a gene product’s functionally annotated 3D structure and residue properties. The variant designs of this work ultimately targeted the reduction of functionality encoded within a gene. These designs and their original mutations may be trading the robustness of a system for more simple and higher performing processes. A comprehensive screen of growth and robustness in multiple conditions would shed light on such tradeoffs, but is technically challenging to perform given the vast range of potential growth conditions and stress combinations *E. coli* is known to have encountered in its evolutionary history.

GlpK’s variant design involved an amino acid substitution that may have increased the glycerol kinase reaction rate. GlpK (glycerol kinase) is part of the pathway for utilizing glycerol as a carbon source. GlpK can form a catalytically active homotetramer and homodimer, though the homotetramer can be allosterically inhibited by fructose-1,6-bisphosphate, a downstream product serving as negative feedback. Mutations to Serine 59 have already been shown to disable homotetramer formation through steric incompatibility^37^. This leaves the homodimer formation, which may accomplish a higher overall rate of glycerol metabolism due to the lack of inhibition.

Mutations to *cyaA* and *crr* may be maintaining Carbon Catabolite Repression (CCR). CyaA (adenylate cyclase) is part of the pathway that generates the activated CRP complex (cAMP-CRP), which goes on to activate genes for multiple secondary carbon source catabolic systems^38^. CyaA’s regulatory region is thought to activate the enzyme^39^. For the condition of glycerol as a carbon source, CyaA would become activated and produce cAMP, therefore activating these catabolic systems^39^. A truncation of *cyaA* has been shown to prevent cAMP production^40^ and the downstream activation of CRP is expected to be nullified. Crr can also play a role in CCR. Unphosphorylated Crr binds to and inhibits GlpK *in vitro*^*38*^, though in the case of a glycerol as a carbon source, phosphorylated Crr should be more abundant. Crr interacts with CyaA and will activate CyaA’s cAMP production if phosphorylated^38^. Though this work does not include CyaA interface data for Crr, the phosphoryl group active sites are located in the middle of the multi-interface residues shared with GlpK; mutations to this interface surface may prevent the interaction with CyaA that activates cAMP production. The repression of these secondary carbon source catabolic systems is known as CCR and is normally enforced by the phosphotransferase system in the presence of glucose^38^. With glycerol as a carbon source, ALE-derived strains may have found a way to maintain CCR through mutations to cyaA while still allowing for the activation of glycerol metabolism. CCR maintenance may ultimately enable a more efficient cell metabolism, since it represses the activation of multiple unnecessary catabolic systems.

As suggested with the *glpK* and *cyaA* ALE mutations from the glycerol carbon-source case study, designs could also result from a combined set of compatible variants, where each variant optimizes for a separate stress. A strain’s environment is naturally composed of multiple conditions, therefore compatible optimizations would be valuable in addressing multi-stress circumstances such as those of industrial scale fermentation (physical, chemical, and biological)^41–43^.

The ALE and designed mutant strains of this work were screened for their phenotypes. The mutant strains were found to be more fit than wild-type in the conditions that initially selected for their presence in ALEs. The ability of the strains hosting designed variants to achieve similar growth rates to beneficial ALE mutants demonstrates the possibility to design variants derived from aggregated ALE data. The similarity between ALE and designed mutant growth rates also suggests that ALE processes of the current scale and time-span represented in this work may not find all possible beneficial sequence changes. This lack of full coverage for beneficial sequence changes by ALE processes may be due to the probability of specific mutational sequence changes. For example, GlpK’s ALE mutation involved a single base pair substitution, while both its designed variants involved two base pair substitutions within the same codon. Designed variant sequence changes may be less likely to occur with ALE. Thus, there likely exists potential for beneficial variants not revealed through ALE, but can be understood through utilization of methods outlined in this study such as the mutation clustering and structural biology approaches.

In the cases involving truncations, ALE mutations and designed strains could produce more benefit than full gene truncations depending on the stressor. This was demonstrated with the *cyaA* mutants, where partial truncations had greater benefit than full truncations with the Δ*pgi* stressor, though no substantial difference in benefit was observed between partial and full truncations of *cyaA* with glycerol as a carbon source. The designs and mutations to *pykF* also involved truncations. Full truncations of *pykF* are thought to be valuable with scarce glucose^44^, where partial truncations may be more valuable in abundant glucose^34^. All of this study’s *pykF* mutations manifested in strains using glucose as a carbon source (Figure 7a) and the majority of the truncations occurred near the middle of the coding sequence, where truncations mostly avoided catalytic sites encoded in the first half of the gene (Figure 7b, Figure 7d). The results from the phenotypic screens demonstrated otherwise: no obvious benefit was gained from *pykF* mutants in competitive glucose growth screens (Figure 7f). There may exist conditions in which a partial truncation to *pykF* is more beneficial than a full truncation, though these conditions are currently unclear with this study’s analysis.

The methods in this work demonstrate the value in leveraging diverse public resources to describe ALE variants, but are not without limitations. The success of adequately characterizing the beneficial mechanisms of mutations relies upon the availability of genome annotations local to the mutations of interest as well as tools that can describe the nature and magnitude of a mutation’s severity. The efforts of this work leveraged an already available whole-genome multi-scale annotation framework and further enriched mutation annotations local to features genes of interest. These additional annotations, coming from resources such as Uniprot^45^, EcoCyc^46^, Pfam^47^, and Mutfunc^48^, are available for the whole genome, though efforts have not yet been made to consolidate them into a unified computational resource. Additionally, specific mutation characterization greatly benefitted from investigating the other mutations local to a feature of interest. Variants can therefore serve as an additional genome annotations, and are especially useful if described by the conditions they were found in. There also exists the potential to grow this set of consolidated mutations with those from other studies and with natural variants. Traditional genome annotations typically don’t include variants, though recent efforts have developed a bioinformatic resource that combines genome annotations as well as variants: the Bitome^49^. The Bitome could serve as the locus for consolidating whole-genome multi-scale feature annotations along with variants and their metadata for a comprehensive, high-resolution, genomic resource per organism.

## 4. Conclusion

This work demonstrates how to design strains from aggregated ALE mutational data. Meta-analysis methods involving nucleotide-level mutation data, rich functional annotations, and predicted mutation effects were used to anticipate the general characteristics of ALE-data-driven strain designs. These predictions described condition-specific designs involving only a small number sequence changes that primarily target genes with both truncating and non-truncating effects. A workflow was developed that executes the meta-analysis and structural biology methods in an order that derives specific nucleotide-level sequence variants applicable to specific conditions from the broader aggregated ALE dataset. Two case-studies were included to serve as proof-of-concept for ALE-data-driven strain designs and may hold value for applications in microbial cell factories and beyond. The case-studies demonstrated beneficial designs based on point mutations and partial-truncations, where mutation functional targets were highlighted by either the accumulation of truncated codons on the gene sequence or point mutation clustering on the 3D structure. In the case with point mutations, the predicted effects of mutations on gene-product properties were necessary to elucidate ALE mutation objectives. Designs were also shown to be beneficial for multiple stressors and there exists mutational evidence that designs can be combined in the same strain. Finally, depending on the stressor, partial truncations were shown to be more beneficial than full gene knockouts. Together, these results demonstrate how aggregated public ALE data and data-driven strain design methods can reveal nucleotide-level design variables currently inaccessible to rational design methods. Until rational methods can predict all possible biological paths between genotypes and phenotypes, data-driven methods will continue to provide value towards strain design.

## Methods

### Biological material

All mutants used the base strain of *E. coli* K-12 MG1655 (ATCC 47076). Mutant strains were generated following a CRISPR/Cas9-assisted protocol outlined by Zhao *et al*.^*50*^ (Supplementary Table 10). This method relies on Cas9 to cut the genome of the starting strain while leaving intact the successfully mutated genome. A single plasmid encoding the CRISPR/Cas9 and Lambda Red Recombinase systems along with repair arms and a 20 nt guide RNA targeting the starting strain sequence was constructed using Golden Gate Assembly. In this case the repair arms were generated using PCR instead of annealing two oligos. Depending on the target mutation, one of two strategies for the placement of the 20 nucleotide guide RNA were used: 1) spanning the bases targeted for mutation if that region was next to a PAM sequence or 2) close to the region being mutated and next to a PAM sequence that could be eliminated by introducing a synonymous codon with the repair arms. When option 2 was used to introduce a single nucleotide change, one of the 20 bases of the guide RNA unrelated to that position was deliberately changed so that the successfully mutated strain would have two mismatched bases with respect to the guide RNA while the starting strain would have just one mismatch. Colonies were screened using ARMS PCR in which one of the primers was designed to work on the starting strain and not on the mutated sequence. All mutations were verified by Sanger sequencing an amplicon generated with primers targeting the genome distal to the ends of the repair arms to avoid sequencing the plasmid. Finally the plasmid was eliminated by growth at 37°C, verified by parallel plating on media with and without Kanamycin.

Each chromosomal *cyaA, pgi* and *pykF* deletion was introduced using a temperature-sensitive pGE3 carrying λ-recombinases and MAD7 nuclease (MADzyme™) which sequence obtained from Inscripta Inc. Briefly, *E. coli* MG1655 was transformed with a temperature-sensitive pGE3 (Supplementary Table 8, Supplementary Table 9), modified from pMP11 by replacing Cas9 into MAD7^51^. Lambda recombinases were induced with 0.2% arabinose at 37°C for 45 min. Then, induced cells were transformed with 200 ng gRNA plasmid together with 100 pmol synthetic oligo containing each flanking 45 bp homologous sequence for each gene. Upon transformation, cells were recovered at 30°C for 1 hr 30 min. Then, the cells were transferred to 2 ml of LB containing ampicillin and chloramphenicol and grew overnight at 30°C. Knockout transformants were isolated by plating and validated by colony PCR and Sanger sequencing. Loss of guide RNA plasmid was done by growing confirmed isolates in LB containing ampicillin and 200 μg/L of anhydrotetracycline at 30°C for at least 6 hours. Loss of pGE3 was achieved by propagating at 37°C. Loss of both plasmids was validated by checking the cell’s sensitivity to antibiotics. *cyaA* knockout was performed on Δ*pgi* strain for construction of *pgi* and *cyaA* double knockout strain.

### Physiological characterizations

The growth rates of strain clones were screened by inoculating cells from an overnight culture to a low OD and then sampling the OD_600_ until the stationary phase was reached. A linear regression of the log-linear region was computed using the *linregress* function from the *scipy.stats* Python package and the growth rates were determined from the resulting slopes. Screen condition details are given in Supplementary Table 7.

### Residue properties

Residue properties were acquired from the software package *ssbio* ^52^.

### SIFT and ΔΔG scores for the mutation effect prediction of amino acid substitutions

In the case studies, two more effect types were predicted for mutations: deleterious and structural destabilization. Both of these were specific to coding regions. Deleterious effects were assumed according to significant SIFT scores (SIFT score < 0.05)^53^. Structural destabilization was assumed according to predicted significant ΔΔG scores (ΔΔG > 2)^54^. SIFT and ΔΔG scores were acquired from *Mutfunc*^*48*^.

### Gene product feature annotations

GlpK features were acquired from Uniprot^45^, EcoCyc^46^, Pfam^47^, Mutfunc^48^ and individual publications^55^.

### Gene product structures and distances between mutated residues and features

Distances between mutated residues and the residues of functional features were calculated using gene product 3D structures and the Cartesian distance formula. GlpK was represented by the 3EZW PDB model. Structures for Crr, CyaA, and PykF were obtained from those provided with the iML1515 model of *E. coli* K-12 MG1655 ^36^.

### Software scripts

The software scripts supporting the conclusions of this article are available in the following open-access archive repository: https://doi.org/10.5281/zenodo.5108959

### Availability of data and materials

The datasets supporting the conclusions of this article are available in the following open-access archive repository: https://doi.org/10.5281/zenodo.5108959 These datasets are also available in the ALEdb database^22^.

### Mutation data cleaning

The mutations from ALEdb are initially described relative to the sample and the genome reference. Since some of these experiments include midpoint samples, mutations that emerge within a midpoint sample and carried through to an endpoint sample can get counted more than once per ALE, which would result in an inappropriately inflated mutation count. Unique ALE mutations were therefore only considered once per ALE. Starting strain mutations were filtered out of the ALE experiment mutation datasets according to their publications after being exported from ALEdb. ALE replicates containing hypermutators were also removed from the dataset. An ALE replicate was predicted to host a hypermutator strain if two conditions were met: 1) a hypermutator gene^56^ was mutated within the ALE replicate, and 2) the number of mutations found within the ALE replicate was labelled as an outlier relative to boxplot quartiles when considering the distribution of mutation counts for all ALE replicates used in this study. Mutations in population samples with a frequency below 50% were filtered out to instead focus on mutations that demonstrate dominant selection within a sample. In calculating correlations between mutated genomic features and generating the network diagrams of multi-scaled mutated features, large deletions were removed to filter out large sets of mutated features that were only mutated once.

### Quantitative plots

Unless otherwise stated, figure plots were generated using *Matplotlib* version 3.0.3^57^ and *Seaborn* version 0.11^58^ or Plotly^59^ Python software packages.

### Network diagrams

The network diagrams of multiscale mutated features were generated using Cytoscape.js^60^.

### Mutation Needle Plots

The mutation needle plots were generated using the *trackViewer* R software package^61^.

### Oncoplots

The oncoplots were generated using the ComplexHeatmap R software package^62^.

### 3D protein structures

The visualizations for the 3D protein structures were generated using the NGL software package^63^.

## Abbreviations

ALE: adaptive laboratory evolution
SNP: single nucleotide polymorphism
DEL: deletion
MOB: mobile insertion elements
INS: insertion
SUB: substitution
AMP: amplification
TFBS: transcription factor binding site
RBS: ribosomal binding site
SV: structural variant
SIFT: Sorting Intolerant from Tolerant
PykF: Pyruvate kinase I
GlpK: glycerol kinase
CyaA: Adenylate cyclase
Crr: PTS system glucose-specific EIIA component
PTS: phosphotransferase system
EIIA: Enzyme II A
CCR: carbon catabolite repression
cAMP-CRP: activated CRP complex
CRP: cAMP receptor protein
cAMP: cyclic AMP
ΔΔG: The predicted difference between the free energy of unfolding the protein structure before and after the variant.

## Author information

### Author contribution

Conceptualization: PVP, AMF, BOP

Conceptual support: DCZ, JTY, AMF, BOP, JJ

Technical support: MW, SS, CD, JJ, RS, LY, SHK, EO

Writing: PVP, DCZ, JTY, AMF, BOP

## Acknowledgments

The authors gratefully acknowledge Robin Cai and Douglas McCloskey for their help with the technologies involved in this project and Marc Abrams for his editorial comments. The authors gratefully acknowledge Connor Olson for his support with the characterization experiment work.

## Supporting Information

Supplementary Document: Supporting figures.

Supplementary Table 1: Describes the mutation count for each genomic feature associated with glycerol as a carbon source.

Supplementary Table 2: Describes the mutation count for each operon associated with glycerol as a carbon source.

Supplementary Table 3: Describes the mutation count for each regulon associated with glycerol as a carbon source.

Supplementary Table 4: Describes the mutation count for each genomic feature associated with toxic concentrations of isobutyric acid.

Supplementary Table 5: Describes the mutation count for each operon associated with toxic concentrations of isobutyric acid.

Supplementary Table 6: Describes the mutation count for each pathway associated with toxic concentrations of isobutyric acid.

Supplementary Table 7: Describes the mutation count for each regulon associated with toxic concentrations of isobutyric acid.

## References

(1) García-Granados, R., Lerma-Escalera, J. A., Morones-Ramírez, J. R. Metabolic Engineering and Synthetic Biology: Synergies, Future, and Challenges. Front Bioeng Biotechnol 2019, 7, 36.

(2) Nielsen, J., Keasling, J. D. Engineering Cellular Metabolism. Cell 2016, 164 (6), 1185–1197.

(3) Lee, S. Y., Kim, H. U. Systems Strategies for Developing Industrial Microbial Strains. Nat. Biotechnol. 2015, 33 (10), 1061–1072.

(4) Sandberg, T. E., Salazar, M. J., Weng, L. L., Palsson, B. O., Feist, A. M. The Emergence of Adaptive Laboratory Evolution as an Efficient Tool for Biological Discovery and Industrial Biotechnology. Metab. Eng. 2019. https://doi.org/10.1016/j.ymben.2019.08.004.

(5) Jantama, K., Haupt, M. J., Svoronos, S. A., Zhang, X., Moore, J. C., Shanmugam, K. T., Ingram, L. O. Combining Metabolic Engineering and Metabolic Evolution to Develop Nonrecombinant Strains of Escherichia Coli C That Produce Succinate and Malate. Biotechnol. Bioeng. 2008, 99 (5), 1140–1153.

(6) Luo, H., Hansen, A. S. L., Yang, L., Schneider, K., Kristensen, M., Christensen, U., Christensen, H. B., Du, B., Özdemir, E., Feist, A. M., Keasling, J. D., Jensen, M. K., Herrgård, M. J., Palsson, B. O. Coupling S-Adenosylmethionine-Dependent Methylation to Growth: Design and Uses. PLoS Biol. 2019, 17 (3), e2007050.

(7) McCloskey, D., Xu, S., Sandberg, T. E., Brunk, E., Hefner, Y., Szubin, R., Feist, A. M., Palsson, B. O. Evolution of Gene Knockout Strains of E. Coli Reveal Regulatory Architectures Governed by Metabolism. Nat. Commun. 2018, 9 (1), 3796.

(8) Choe, D., Lee, J. H., Yoo, M., Hwang, S., Sung, B. H., Cho, S., Palsson, B., Kim, S. C., Cho, B.-K. Adaptive Laboratory Evolution of a Genome-Reduced Escherichia Coli. Nat. Commun. 2019, 10 (1), 935.

(9) Wannier, T. M., Kunjapur, A. M., Rice, D. P., McDonald, M. J., Desai, M. M., Church, G. M. Adaptive Evolution of Genomically Recoded Escherichia Coli. Proc. Natl. Acad. Sci. U. S. A. 2018, 115 (12), 3090–3095.

(10) Zelle, R. M., Harrison, J. C., Pronk, J. T., van Maris, A. J. A. Anaplerotic Role for Cytosolic Malic Enzyme in Engineered Saccharomyces Cerevisiae Strains. Appl. Environ. Microbiol. 2011, 77 (3), 732–738.

(11) Carroll, S. M., Marx, C. J. Evolution after Introduction of a Novel Metabolic Pathway Consistently Leads to Restoration of Wild-Type Physiology. PLoS Genet. 2013, 9 (4), e1003427.

(12) Qi, Y., Liu, H., Chen, X., Liu, L. Engineering Microbial Membranes to Increase Stress Tolerance of Industrial Strains. Metab. Eng. 2019, 53, 24–34.

(13) Wang, S., Sun, X., Yuan, Q. Strategies for Enhancing Microbial Tolerance to Inhibitors for Biofuel Production: A Review. Bioresour. Technol. 2018, 258, 302–309.

(14) Bellissimi, E., van Dijken, J. P., Pronk, J. T., van Maris, A. J. A. Effects of Acetic Acid on the Kinetics of Xylose Fermentation by an Engineered, Xylose-Isomerase-Based Saccharomyces Cerevisiae Strain. FEMS Yeast Res. 2009, 9 (3), 358–364.

(15) Heer, D., Sauer, U. Identification of Furfural as a Key Toxin in Lignocellulosic Hydrolysates and Evolution of a Tolerant Yeast Strain. Microb. Biotechnol. 2008, 1 (6), 497–506.

(16) Rajaraman, E., Agrawal, A., Crigler, J., Seipelt-Thiemann, R., Altman, E., Eiteman, M. A. Transcriptional Analysis and Adaptive Evolution of Escherichia Coli Strains Growing on Acetate. Appl. Microbiol. Biotechnol. 2016, 100 (17), 7777–7785.

(17) Lee, D.-H., Palsson, B. Ø. Adaptive Evolution of Escherichia Coli K-12 MG1655 during Growth on a Nonnative Carbon Source, L-1,2-Propanediol. Appl. Environ. Microbiol. 2010, 76 (13), 4158–4168.

(18) Guzmán, G. I., Sandberg, T. E., LaCroix, R. A., Nyerges, Á., Papp, H., de Raad, M., King, Z. A., Hefner, Y., Northen, T. R., Notebaart, R. A., Pál, C., Palsson, B. O., Papp, B., Feist, A. M. Enzyme Promiscuity Shapes Adaptation to Novel Growth Substrates. Mol. Syst. Biol. 2019, 15 (4), e8462.

(19) Notebaart, R. A., Kintses, B., Feist, A. M., Papp, B. Underground Metabolism: Network-Level Perspective and Biotechnological Potential. Curr. Opin. Biotechnol. 2018, 49, 108–114.

(20) Van den Bergh, B., Swings, T., Fauvart, M., Michiels, J. Experimental Design, Population Dynamics, and Diversity in Microbial Experimental Evolution. Microbiol. Mol. Biol. Rev. 2018, 82 (3). https://doi.org/10.1128/MMBR.00008-18.

(21) Wang, X., Zorraquino, V., Kim, M., Tsoukalas, A., Tagkopoulos, I. Predicting the Evolution of Escherichia Coli by a Data-Driven Approach. Nat. Commun. 2018, 9 (1), 3562.

(22) Phaneuf, P. V., Gosting, D., Palsson, B. O., Feist, A. M. ALEdb 1.0: A Database of Mutations from Adaptive Laboratory Evolution Experimentation. Nucleic Acids Res. 2019, 47 (D1), D1164–D1171.

(23) Phaneuf, P. V., Yurkovich, J. T., Heckmann, D., Wu, M., Sandberg, T. E., King, Z. A., Tan, J., Palsson, B. O., Feist, A. M. Causal Mutations from Adaptive Laboratory Evolution Are Outlined by Multiple Scales of Genome Annotations and Condition-Specificity. BMC Genomics 2020, 21 (1), 514.

(24) Rudrapatna, V. A., Butte, A. J. Open Data Informatics and Data Repurposing for IBD. Nat. Rev. Gastroenterol. Hepatol. 2018, 15 (12), 715–716.

(25) Stransky, N., Cerami, E., Schalm, S., Kim, J. L., Lengauer, C. The Landscape of Kinase Fusions in Cancer. Nat. Commun. 2014, 5, 4846.

(26) Kavvas, E. S., Catoiu, E., Mih, N., Yurkovich, J. T., Seif, Y., Dillon, N., Heckmann, D., Anand, A., Yang, L., Nizet, V., Monk, J. M., Palsson, B. O. Machine Learning and Structural Analysis of Mycobacterium Tuberculosis Pan-Genome Identifies Genetic Signatures of Antibiotic Resistance. Nat. Commun. 2018, 9 (1), 4306.

(27) Bailey, M. H., Tokheim, C., Porta-Pardo, E., Sengupta, S., Bertrand, D., Weerasinghe, A., Colaprico, A., Wendl, M. C., Kim, J., Reardon, B., Ng, P. K.-S., Jeong, K. J., Cao, S., Wang, Z., Gao, J., Gao, Q., Wang, F., Liu, E. M., Mularoni, L., Rubio-Perez, C., Nagarajan, N., Cortés-Ciriano, I., Zhou, D. C., Liang, W.-W., Hess, J. M., Yellapantula, V. D., Tamborero, D., Gonzalez-Perez, A., Suphavilai, C., Ko, J. Y., Khurana, E., Park, P. J., Van Allen, E. M., Liang, H., MC3 Working Group; Cancer Genome Atlas Research Network; Lawrence, M. S., Godzik, A., Lopez-Bigas, N., Stuart, J., Wheeler, D., Getz, G., Chen, K., Lazar, A. J., Mills, G. B., Karchin, R., Ding, L. Comprehensive Characterization of Cancer Driver Genes and Mutations. Cell 2018, 173 (2), 371–385.e18.

(28) Zielinski, D. C., Patel, A., Palsson, B. O. The Expanding Computational Toolbox for Engineering Microbial Phenotypes at the Genome Scale. Microorganisms 2020, 8 (12), 2050.

(29) Clomburg, J. M., Gonzalez, R. Biofuel Production in Escherichia Coli: The Role of Metabolic Engineering and Synthetic Biology. Appl. Microbiol. Biotechnol. 2010, 86 (2), 419–434.

(30) Shams Yazdani, S., Gonzalez, R. Engineering Escherichia Coli for the Efficient Conversion of Glycerol to Ethanol and Co-Products. Metab. Eng. 2008, 10 (6), 340–351.

(31) Willrodt, C., David, C., Cornelissen, S., Bühler, B., Julsing, M. K., Schmid, A. Engineering the Productivity of Recombinant Escherichia Coli for Limonene Formation from Glycerol in Minimal Media. Biotechnol. J. 2014, 9 (8), 1000–1012.

(32) Lennen, R. M., Jensen, K., Mohammed, E. T., Malla, S., Börner, R. A., Chekina, K., Özdemir, E., Bonde, I., Koza, A., Maury, J., Pedersen, L. E., Schöning, L. Y., Sonnenschein, N., Palsson, B. O., Sommer, M. O. A., Feist, A. M., Nielsen, A. T., Herrgård, M. J. Adaptive Laboratory Evolution Reveals General and Specific Chemical Tolerance Mechanisms and Enhances Biochemical Production. bioRxiv, 2019, 634105. https://doi.org/10.1101/634105.

(33) Zhang, K., Woodruff, A. P., Xiong, M., Zhou, J., Dhande, Y. K. A Synthetic Metabolic Pathway for Production of the Platform Chemical Isobutyric Acid. ChemSusChem 2011, 4 (8), 1068–1070.

(34) Sandberg, T. E., Pedersen, M., LaCroix, R. A., Ebrahim, A., Bonde, M., Herrgard, M. J., Palsson, B. O., Sommer, M., Feist, A. M. Evolution of Escherichia Coli to 42 °C and Subsequent Genetic Engineering Reveals Adaptive Mechanisms and Novel Mutations. Mol. Biol. Evol. 2014, 31 (10), 2647–2662.

(35) Sandberg, T. E., Lloyd, C. J., Palsson, B. O., Feist, A. M. Laboratory Evolution to Alternating Substrate Environments Yields Distinct Phenotypic and Genetic Adaptive Strategies. Appl. Environ. Microbiol. 2017, 83 (13). https://doi.org/10.1128/AEM.00410-17.

(36) Monk, J. M., Lloyd, C. J., Brunk, E., Mih, N., Sastry, A., King, Z., Takeuchi, R., Nomura, W., Zhang, Z., Mori, H., Feist, A. M., Palsson, B. O. iML1515, a Knowledgebase That Computes Escherichia Coli Traits. Nat. Biotechnol. 2017, 35 (10), 904–908.

(37) Bystrom, C. E., Pettigrew, D. W., Branchaud, B. P., O’Brien, P., Remington, S. J. Crystal Structures of Escherichia Coli Glycerol Kinase Variant S58-->W in Complex with Nonhydrolyzable ATP Analogues Reveal a Putative Active Conformation of the Enzyme as a Result of Domain Motion. Biochemistry 1999, 38 (12), 3508–3518.

(38) Görke, B., Stülke, J. Carbon Catabolite Repression in Bacteria: Many Ways to Make the Most out of Nutrients. Nat. Rev. Microbiol. 2008, 6 (8), 613–624.

(39) Saier, M. H., Jr; Kukita, C., Zhang, Z. Transposon-Mediated Directed Mutation in Bacteria and Eukaryotes. Front. Biosci. 2017, 22, 1458–1468.

(40) McCloskey, D., Xu, S., Sandberg, T. E., Brunk, E., Hefner, Y., Szubin, R., Feist, A. M., Palsson, B. O. Adaptive Laboratory Evolution Resolves Energy Depletion to Maintain High Aromatic Metabolite Phenotypes in Escherichia Coli Strains Lacking the Phosphotransferase System. Metab. Eng. 2018, 48, 233–242.

(41) Wehrs, M., Tanjore, D., Eng, T., Lievense, J., Pray, T. R., Mukhopadhyay, A. Engineering Robust Production Microbes for Large-Scale Cultivation. Trends Microbiol. 2019, 27 (6), 524–537.

(42) Lennen, R. M., Herrgård, M. J. Combinatorial Strategies for Improving Multiple-Stress Resistance in Industrially Relevant Escherichia Coli Strains. Appl. Environ. Microbiol. 2014, 80 (19), 6223–6242.

(43) Qiao, W., Qiao, Y., Liu, F., Zhang, Y., Li, R., Wu, Z., Xu, H., Saris, P. E. J., Qiao, M. Engineering Lactococcus Lactis as a Multi-Stress Tolerant Biosynthetic Chassis by Deleting the Prophage-Related Fragment. Microb. Cell Fact. 2020, 19 (1), 225.

(44) Barrick, J. E., Yu, D. S., Yoon, S. H., Jeong, H., Oh, T. K., Schneider, D., Lenski, R. E., Kim, J. F. Genome Evolution and Adaptation in a Long-Term Experiment with Escherichia Coli. Nature 2009, 461 (7268), 1243–1247.

(45) UniProt Consortium. UniProt: A Worldwide Hub of Protein Knowledge. Nucleic Acids Res. 2019, 47 (D1), D506–D515.

(46) Keseler, I. M., Mackie, A., Santos-Zavaleta, A., Billington, R., Bonavides-Martínez, C., Caspi, R., Fulcher, C., Gama-Castro, S., Kothari, A., Krummenacker, M., Latendresse, M., Muñiz-Rascado, L., Ong, Q., Paley, S., Peralta-Gil, M., Subhraveti, P., Velázquez-Ramírez, D. A., Weaver, D., Collado-Vides, J., Paulsen, I., Karp, P. D. The EcoCyc Database: Reflecting New Knowledge about Escherichia Coli K-12. Nucleic Acids Res. 2017, 45 (D1), D543–D550.

(47) El-Gebali, S., Mistry, J., Bateman, A., Eddy, S. R., Luciani, A., Potter, S. C., Qureshi, M., Richardson, L. J., Salazar, G. A., Smart, A., Sonnhammer, E. L. L., Hirsh, L., Paladin, L., Piovesan, D., Tosatto, S. C. E., Finn, R. D. The Pfam Protein Families Database in 2019. Nucleic Acids Res. 2019, 47 (D1), D427–D432.

(48) Wagih, O., Galardini, M., Busby, B. P., Memon, D., Typas, A., Beltrao, P. A Resource of Variant Effect Predictions of Single Nucleotide Variants in Model Organisms. Mol. Syst. Biol. 2018, 14 (12), e8430.

(49) Lamoureux, C. R., Choudhary, K. S., King, Z. A., Sandberg, T. E., Gao, Y., Sastry, A. V., Phaneuf, P. V., Choe, D., Cho, B.-K., Palsson, B. O. The Bitome: Digitized Genomic Features Reveal Fundamental Genome Organization. Nucleic Acids Res. 2020, 48 (18), 10157–10163.

(50) Zhao, D., Yuan, S., Xiong, B., Sun, H., Ye, L., Li, J., Zhang, X., Bi, C. Development of a Fast and Easy Method for Escherichia Coli Genome Editing with CRISPR/Cas9. Microb. Cell Fact. 2016, 15 (1), 205.

(51) Mehrer, C. R., Incha, M. R., Politz, M. C., Pfleger, B. F. Anaerobic Production of Medium-Chain Fatty Alcohols via a β-Reduction Pathway. Metab. Eng. 2018, 48, 63–71.

(52) Mih, N., Brunk, E., Chen, K., Catoiu, E., Sastry, A., Kavvas, E., Monk, J. M., Zhang, Z., Palsson, B. O. Ssbio: A Python Framework for Structural Systems Biology. Bioinformatics 2018, 34 (12), 2155–2157.

(53) Vaser, R., Adusumalli, S., Leng, S. N., Sikic, M., Ng, P. C. SIFT Missense Predictions for Genomes. Nat. Protoc. 2016, 11 (1), 1–9.

(54) Mosca, R., Céol, A., Aloy, P. Interactome3D: Adding Structural Details to Protein Networks. Nat. Methods 2013, 10 (1), 47–53.

(55) Eppler, T., Postma, P., Schütz, A., Völker, U., Boos, W. Glycerol-3-Phosphate-Induced Catabolite Repression in Escherichia Coli. J. Bacteriol. 2002, 184 (11), 3044–3052.

(56) Horst, J. P., Wu, T. H., Marinus, M. G. Escherichia Coli Mutator Genes. Trends Microbiol. 1999, 7 (1), 29–36.

(57) Hunter, J. D. Matplotlib: A 2D Graphics Environment. Computing in Science Engineering 2007, 9 (3), 90–95.

(58) Waskom, M., Gelbart, M., Botvinnik, O., Ostblom, J., Hobson, P., Lukauskas, S., Gemperline, D. C., Augspurger, T., Halchenko, Y., Warmenhoven, J., Cole, J. B., de Ruiter, J., Vanderplas, J., Hoyer, S., Pye, C., Miles, A., Swain, C., Meyer, K., Martin, M., Bachant, P., Quintero, E., Kunter, G., Villalba, S. Brian; Fitzgerald, C., Evans, C., Williams, M. L., O’Kane, D., Yarkoni, T., Brunner, T. Mwaskom/seaborn: v0.11.1 (December 2020); 2020. https://doi.org/10.5281/zenodo.4379347.

(59) Inc, P. T. Collaborative Data Science with Plotly. Montréal, QC 2015.

(60) Franz, M., Lopes, C. T., Huck, G., Dong, Y., Sumer, O., Bader, G. D. Cytoscape.js: A Graph Theory Library for Visualisation and Analysis. Bioinformatics 2016, 32 (2), 309–311.

(61) Ou, J., Zhu, L. J. trackViewer: A Bioconductor Package for Interactive and Integrative Visualization of Multi-Omics Data. Nat. Methods 2019, 16 (6), 453–454.

(62) Gu, Z., Eils, R., Schlesner, M. Complex Heatmaps Reveal Patterns and Correlations in Multidimensional Genomic Data. Bioinformatics 2016, 32 (18), 2847–2849.

(63) Rose, A. S., Bradley, A. R., Valasatava, Y., Duarte, J. M., Prlic, A., Rose, P. W. NGL Viewer: Web-Based Molecular Graphics for Large Complexes. Bioinformatics 2018, 34 (21), 3755–3758.

